# Decay drives RNA abundance regulation using three distinct regulatory mechanisms

**DOI:** 10.1101/2025.05.09.653099

**Authors:** Reed S. Sorenson, Leslie E. Sieburth

## Abstract

RNA decay is essential for maintenance of normal RNA abundances; however how RNA decay is regulated to contribute to changes in RNA abundances is poorly understood. Here, we addressed this question by analyzing rates of RNA abundance change, RNA half-lives (*t*1/2s), and transcription rates in stimulated Arabidopsis leaf cells. This revealed three mechanisms by which decay influenced RNA abundance changes. First, the biggest changes in RNA abundances resulted from *t*1/2 changes that reinforced transcriptional regulation (synergistic). Modest RNA abundance changes arose from a second mechanism in which *t*1/2 changes opposed transcriptional regulation (oppositional). Finally, RNA decay alone also contributed to RNA abundance change, and RNA decay’s measured capacity influence RNA abundances was similar to that of transcription. RNA decay also contributed to transcriptome homeostasis through stimulus-induced RNA buffering. Oppositional and buffering regulation shared key features, including excessive and commensurate rate changes, which suggested use of a shared regulatory mechanism which we call countercyclical regulation. In this study, countercyclical regulation was widespread and used for regulation of 90% of the RNAs with *t*1/2 regulation.

## INTRODUCTION

Cells respond to developmental and environmental perturbations in part by altering messenger RNA (mRNA) abundances. The identity of responsive RNAs can be highly informative; they can indicate developmental status or perception of environmental stress, and can be used to test hypotheses, for example in working out gene regulatory networks. However, despite RNA abundances being governed by just two processes, transcription and decay, we know relatively little about its regulation by RNA decay. An implicit assumption is that mRNA abundances go up because transcription rates (TRs) increase and perhaps RNA decay rates (*t*1/2s) diminish, and that mRNA abundances go down because TRs diminish and perhaps RNA decay rates increase. However, whether transcription and decay work together in this way, and how frequently RNA stabilities are regulated to contribute to RNA abundance regulation, is largely unknown.

Decay of RNA is a highly regulated process; it initiates by removal of 5’ and 3’ structures that confer stability (5’ cap and 3’ tail) or through internal cleavage. The remaining terminal nucleotides are then vulnerable to exoribonucleases such as XRN1/XRN4^1,2^, SOV^3^, or the RNA exosome^4^. RNA stability is regulated by a variety of mechanisms, e.g., signal transduction can activate decay machinery by phosphorylation^5,6^; DNE1 can initiate decay by internal cleavage^7–9^; and RNA binding proteins or RNA modification can influence recruitment of decay machinery^10–14^. Additionally, RNA localization such as sequestration in processing bodies or stress granules can affect stability^15–17^. However, despite understanding these mechanisms of regulation, the extent and manner in which RNA decay contributes to changing RNA abundances is still poorly understood.

A few studies have addressed the contributions of RNA decay regulation to gene expression. In animals, 20-30% of RNAs are regulated by the combined influences of transcription and decay, whereas in yeast decay contributes much more widely^18–24^. Studies using plants, though typically focused on small sets of genes, have demonstrated decay-influenced abundance regulation in response to time-of-day^25^, iron-mediated oxidative stress^26^, boron treatment^27^, osmotic stress^6,28^, and temperature^29–32^. However, a systems-level view of transcription and decay interactions focused on RNA abundance regulation is still lacking. A challenge for understanding RNA abundance regulation is that its two regulatory processes, transcription and decay, mostly occur in separate compartments.

Nevertheless, there is evidence that these processes can be coordinated. One example is RNA buffering, which was discovered in a *Saccharomyces cerevisiae* mutant with an RNA polymerase II defect, was found to have normal RNA abundances because of compensatory changes in mRNA decay rates^33^. RNA buffering is now known to be widespread in yeast genotypes with defects in transcription and decay^34,35^. Its mechanism is not fully understood, but in yeast, shuttling of RNA decay enzymes between the nucleus and cytoplasm, especially XRN1, is a central component^34,36–39^. RNA buffering has also been identified in Arabidopsis^40^ and in animal cells^36,37,41,42^. For example, in Arabidopsis, dysfunction of SUPPRESSOR OF VARICOSE (SOV)/DIS3L2 evoked RNA buffering; 42% of the transcriptome underwent significant shifts in decay which contributed to maintenance of normal RNA abundances^40^. However, whether buffering-like coordination between transcription and decay extends to other modes of gene expression, including those leading to differential RNA abundances, remains unknown.

Here, we investigated the roles of RNA decay in abundance regulation using leaf cells as they responded to a developmental stimulus. This stimulus led to widespread changes in RNA *t*1/2s; 22% of the analyzed RNAs had differing mock and stimulus *t*1/2. These *t*1/2 changes affected RNA abundances using three distinct mechanisms. First, RNA *t*1/2 changes worked alongside changes in transcription to bring about large changes in RNA abundances. This mechanism, which we call synergistic regulation, was consistent with implicit assumptions about gene regulation. Second, abundances of some RNAs were regulated via decay (only). Finally, a widely-used mechanism for abundance regulation entailed opposing changes in *t*1/2s and transcription. These rate changes were excessive, sometimes dramatically so, and typically led to modest transcription-driven changes in abundance. We called this mechanism oppositional because of the apparent antagonism between transcription and decay. Similar excessive rate changes were observed for a set of RNAs whose abundances were steady and so appeared to be buffered. Oppositionally regulated and buffered RNAs were only distinguishable by their impact on RNA abundance, suggesting regulation by a common mechanism which we call countercyclical regulation.

## RESULTS

### A dynamic system for study of RNA abundance regulation

To determine how RNA decay contributes to changes in RNA abundances, we used a well-characterized leaf vascular transdifferentiation system^43–46^ in which many RNAs change abundances in response to a stimulus (Supplemental Data File 1). Developmental progression of the stimulus response was monitored by following expression of the preprocambial cell reporter *proAtHB8:4xYFP*^47–49^. *AtHB8* expression is normally activated in narrow cell files that later differentiate into organized leaf veins^50,51^. In stimulus-treated leaves, *proAtHB8:4xYFP* expression had expanded beyond these narrow cell files and by 72 h cells of interveinal regions had thick and lignified cell walls that resembled xylem tracheary elements (Supplemental Figure S1A, S1B, S1C). These observations were consistent with the system’s described developmental progression and demonstrated reliable elevated *proAtHB8:4xYFP* fluorescence after 12 h of stimulus (Figure 1A). Vascular induction was confirmed by qPCR analysis of *AtHB8* and *DOF5.8*, another early vascular transcription factor (Figure 1B), while a later-expressing phloem-specific gene, *SMXL5*, showed peak expression later (between 12 and 16 h)^52–56^. These observations indicated that vascular-related gene expression was ectopically activated and proceeding along a stereotypical timeline. Because the molecular signature of vascular induction was apparent at 8 h while cell morphologies were not obviously altered (Supplemental Figure S1B), this timepoint seemed suitable for analysis of RNA decay’s roles in abundance regulation.

**Figure 1.**
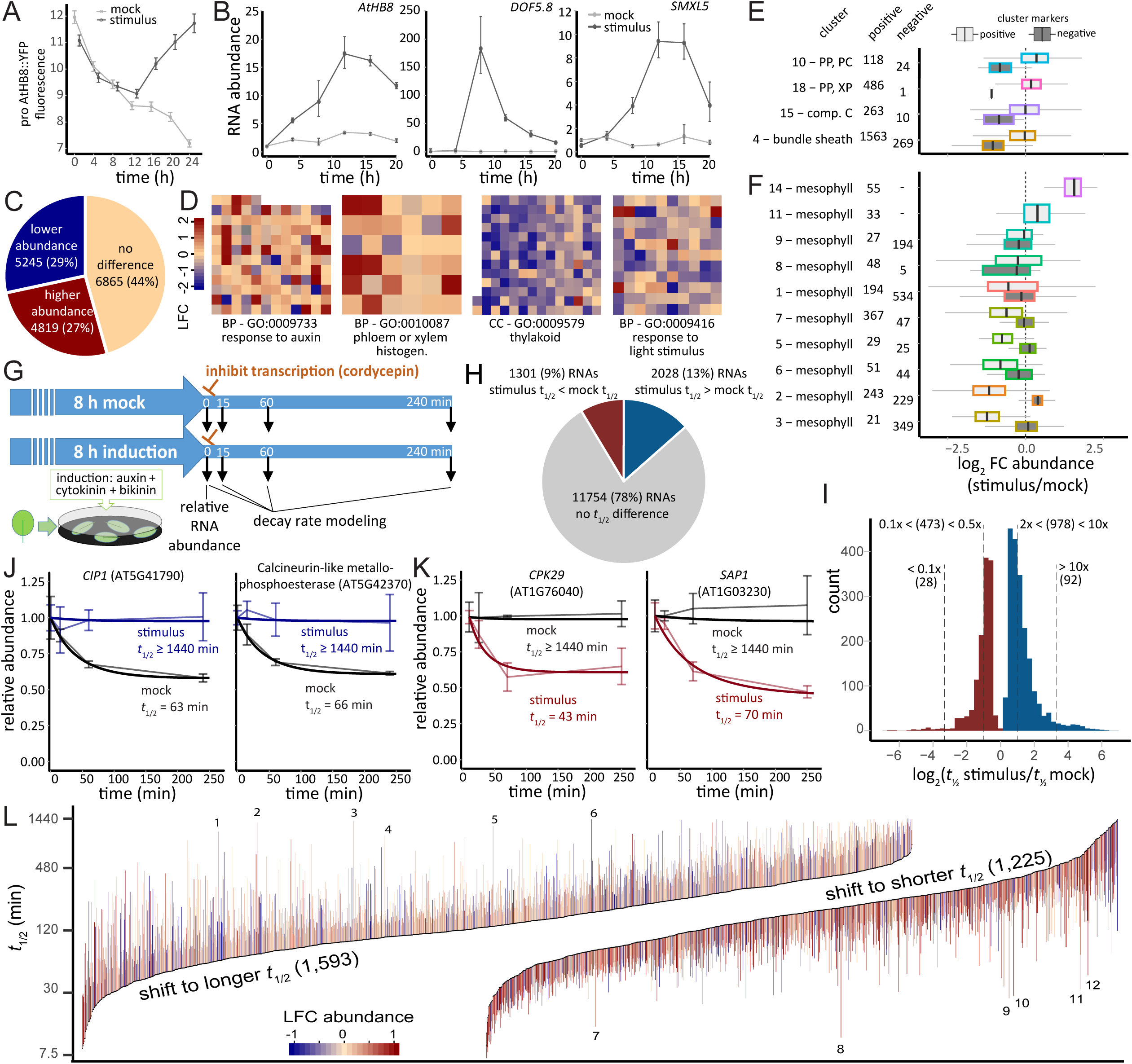
Vascular developmental stimulus induces active cellular reprogramming (A) Effect of stimulus on an early procambial marker. YFP fluorescence was measured as mean pixel intensity x 10^-3^ from 2 mm diameter leaf disks of proAtHB8::4xYFP plants over 24 h treatment. (B) Effect of stimulus on relative RNA abundance of vascular differentiation markers. RNA abundance was measured by qRT-PCR over 20 h of treatment and normalized to the *UBC10* reference RNA. (C) Pie chart showing the proportions of RNAs with increased or decreased RNA abundance (FDR < 0.05) by RNA-seq at 8 h stimulus treatment compared with mock. (D) Effect of stimulus on RNA abundances of members of the *phloem or xylem histogenesis*, *response to auxin*, *thylakoid*, and *response to light stimulus* gene ontologies. Color squares represent log2 relative RNA abundance (LFC) of GO members. BP, biological process; CC, cellular component. (E, F) Stimulus-induced changes in leaf vascular cell-identity (E) and mesophyll (F) markers. Box plot of log2 relative RNA abundance change (8 h stimulus vs mock) of markers identified by leaf single cell RNA-seq clusters generated by Kim et al.^58^. (G) Experimental design schematic for RNA abundance and RNA decay analysis. (H) Pie chart showing proportions of RNAs with increased, decreased, or unchanged *t1/2*s. (I) Distribution of *t*1/2 changes. (J,K) Decay plots showing RNA abundance changes after inhibition of transcription (thin lines, mean±SE) and *t*1/2 estimates based on modeled decay (thick lines) of 8 h stimulus- (blue and red) and mock-treated (black) leaves. (L) Distribution of *t*1/2 (8 h mock, black dots) and shift in *t*1/2 (line segments) for 2,818 RNAs with shifted *t*1/2s in the standard range. Line segment endpoints are positioned at the mock and stimulus *t*1/2s. Line color indicates log2 relative abundance (LFC). See Figure S5A for enlargement of the 100 shortest and longest mock *t*1/2s. Numbered line segments indicate RNAs that have much shorter or longer *t*1/2s in stimulus: (1) Protein kinase family protein with leucine-rich repeat (AT1G35710), (2) TIR-NBS-LRR Disease resistance protein (AT4G36150), (3) CCCH-type zinc finger (AT5G58620), (4) hypothetical protein (AT2G07779), (5) hypothetical protein (AT5G66580), (6) snoRNA (AT1G32385), (7) FTSH11 (AT5G53170), (8) BTB/POZ domain-containing protein (AT5G59140), (9) CID4 (AT3G14010), (10) TRM8 (AT5G26910), (11) transcriptional regulator of RNA PolII, SAGA, subunit (AT4G33890), (12) Disease resistance protein (TIR-NBS-LRR class) (AT4G16900).

To characterize the developmental status of the treated leaves at 8 h, we quantified RNA abundances of mock and stimulus-treated leaf tissue using RNA-seq (Supplemental Data File 1). This yielded relative abundances of 18,152 nuclear-encoded RNAs, 55% of which differed significantly in the mock and stimulus samples (FDR < 0.05) (Figure 1C, Supplemental Figure S1D). These abundance changes resembled early transdifferentiation responses in cotyledons as captured by a microarray time course^46^, and differed from expression of mature vascular tissues of late stimulus responses^45^ (Supplemental Figure S1E-S1F). An early developmental status at 8 h was also supported by gene ontology analysis; stimulus up-regulated RNAs showed enrichment of *response to auxin* (GO: GO:0009733), a stimulus cocktail component, and RNAs of later vascular cell type differentiation not enriched in *phloem or xylem histogenesis* (GO:0010087) (Figure 1D).

Moreover, mesophyll-cell dedifferentiation was supported because stimulus down- regulated RNAs were enriched in the *response to light stimulus* and *thylakoid* GOs (GO:0009416, 73/209; and GO:0009579, 143/148, respectively) (Supplemental Data File 2)^57^.

We also compared 8 h RNA abundances to leaf cell types classified from single-cell RNA sequencing^58^ (see methods, Supplemental Figure S1G, Supplemental Data Files 3 and 4). Stimulus-induced abundance changes included an increase in positive markers for the procambium cell type cluster (no. 10), a decrease in procambium negative markers (Figure 1E), and a strong reduction in negative markers of differentiating and mature vascular cell types (clusters 10, 18, 15, and 4). These changes indicated that cells had initiated vascular differentiation. Cell-type clustering separated mesophyll cells into 10 groups and the 8 h data showed two types of responses. Six clusters showed a reduction of positive markers while negative markers were unchanged or higher (Figure 1F), consistent with the loss of mesophyll identity as the stimulus redirects development toward vascular cell types. Of the remaining four mesophyll cell clusters, positive markers of clusters 11 and 14 were increased as much or more than those of procambium cell cluster 10, suggesting that these mesophyll cell subtypes might be procambial cell precursors. These cell identity-driven shifts include many changes in gene expression, suggesting that the 8 h time point might be a rich system for teasing out roles of RNA decay in RNA abundance regulation.

### Stimulus induces widespread changes in RNA *t*1/2

To determine RNA *t*1/2s, we used the 8 h RNA-seq data described above along with RNA-seq abundances from co-collected tissue treated with the transcription inhibitor cordycepin for 15, 60, or 240 min (Figure 1G). Data were processed through our previously described RNA decay modeling pipeline, which accounts for potential feedback from transcription arrest using a modeling-based approach to estimate initial decay rates. Modeling also provided statistical support for gene-specific *t*1/2 differences in mock and stimulus conditions^40^. We obtained RNA decay rate estimates for 15,083 transcripts. As seen in previous Arabidopsis RNA decay studies, *t*1/2s were spread across 2.5 orders of magnitude^40,59–62^ and *t*1/2s ranged from 5.0 min (*SAUR21* in mock) and 4.6 min (*SAUR14* in stimulus) to ≥ 1,440 min, our capped maximum (Supplemental Data File 1).

We found that 22% of RNAs (3,329) responded to the stimulus by changing RNA *t*1/2 (Figure 1H). These *t*1/2 shifts varied in their magnitude; nearly half (1,571) were greater than 2-fold, and shifts of 120 RNAs were greater than 10-fold (Figure 1I). Some *t*1/2 changes were extraordinarily large. For example, *t*1/2 lengthened 23-fold for RNAs of *CIP1* (*t*1/2 mock = 63 min, *t*1/2 stimulus = 1440 min) and AT5G42370 (*t*1/2 mock = 63 min, *t*1/2 stimulus = 1440 min) (Figure 1J), and shortened 33-fold for *CPK29* (*t*1/2 mock ≥ 1,440 min, *t*1/2 stimulus = 43 min) and 21-fold for *SAP1* (*t*1/2 mock ≥ 1,440 min, *t1*/2 stimulus = 70 min) (Figure 1K).

To assess trends in RNA *t*1/2 shifts, we separated RNAs into two groups: one group was comprised of 2,818 RNAs with *t*1/2s in the standard range (mock and stimulus *t*1/2 ≤ 1440 min) and a second group had at least one extraordinarily long *t*1/2 in mock or stimulus (511, *t*1/2 ≥ 1440 min). RNAs in the standard *t*1/2 range showed similar shifts to shorter and longer *t*1/2 (shift to shorter: mean fold change 2.2, median 1.9; shifts to longer: mean fold change of 2.0, median of 1.8) (Figure 1L, Supplemental Figure S2A). The smooth curvature of these plots indicated that RNAs across the full range of mock *t*1/2s underwent *t*1/2 shifts in both directions, with exceptions being extremely short-lived RNAs, which tended to shift to longer *t*1/2s (75 of the 100 shortest-lived RNAs made this shift) and extremely long-lived RNAs, which tended to shift to shorter *t*1/2s (87 of the 100 longest-lived RNAs) (Supplemental Figure S2B). These patterns suggested that our data approached the upper and lower limits of Arabidopsis RNA *t*1/2s.

The group of RNAs with at least one extraordinarily long t1/2 included 435 RNAs whose standard mock *t1/2* shifted to ≥ 1,440 min, and 76 RNAs with a mock *t*1/2 ≥ 1,440 min that shifted into the standard *t1/2* range (Figure 1J-1K and Supplemental Figure S2A and S2C). The magnitudes of these t1/2 shifts were much larger than for RNAs in the standard range (mean fold change of 3.9, median of 2.8) and so might be entering or exiting sites of RNA sequestration, e.g., condensates such as stress granules or processing bodies^17,63–65^.

Previously we noted that RNAs with rapid turnover were enriched in GOs termed responsive, e.g., to stress or signaling^40^. This suggested that short-lived RNAs might change abundances by altering their t1/2, e.g., by shifting to longer t1/2s. We addressed this idea by comparing *t*1/2 changes across five quintiles of mock *t*1/2. We found that most t1/2 changes occurred among long-lived RNAs (Supplemental Figure S2D), leaving the purpose for short RNAs *t*1/2s an open question.

### Analytical approaches using an early vascular regulatory network

To determine the impact of RNA decay on RNA abundances, we compared two approaches using a 65-gene subset of the data, an early vascular regulatory network (EVR network) (Supplemental Table 1). We obtained RNA abundance and *t*1/2 data for 57 of the EVR network genes, of which 46 (81%) were differentially expressed (Figure 2A and 2B). We compared these abundance changes to mock and stimulus *t*1/2s to infer regulation (Supplemental Figure S3A). Thirty-nine RNAs had higher 8-h abundances in stimulus, and 30 of these had unchanged *t*1/2s, suggesting that their abundance changes were driven by transcription. The remaining included three RNAs that shifted to longer *t*1/2s (e.g., *LBD4*) and six that shifted to shorter *t*1/2s. The shift to longer *t*1/2 likely contributed to the RNA’s abundance change, but the more common outcome, shifts to shorter *t*1/2 would have worked against their increased abundances, and similar observations were made for RNAs with decreased abundances (Figure 2C, Figure S3A, Supplemental Data File 5). An obvious shortcoming of this approach was that it compared an 8 h *t*1/2 measurement to to an abundance change that could have occurred at any time prior to 8 h.

**Figure 2.**
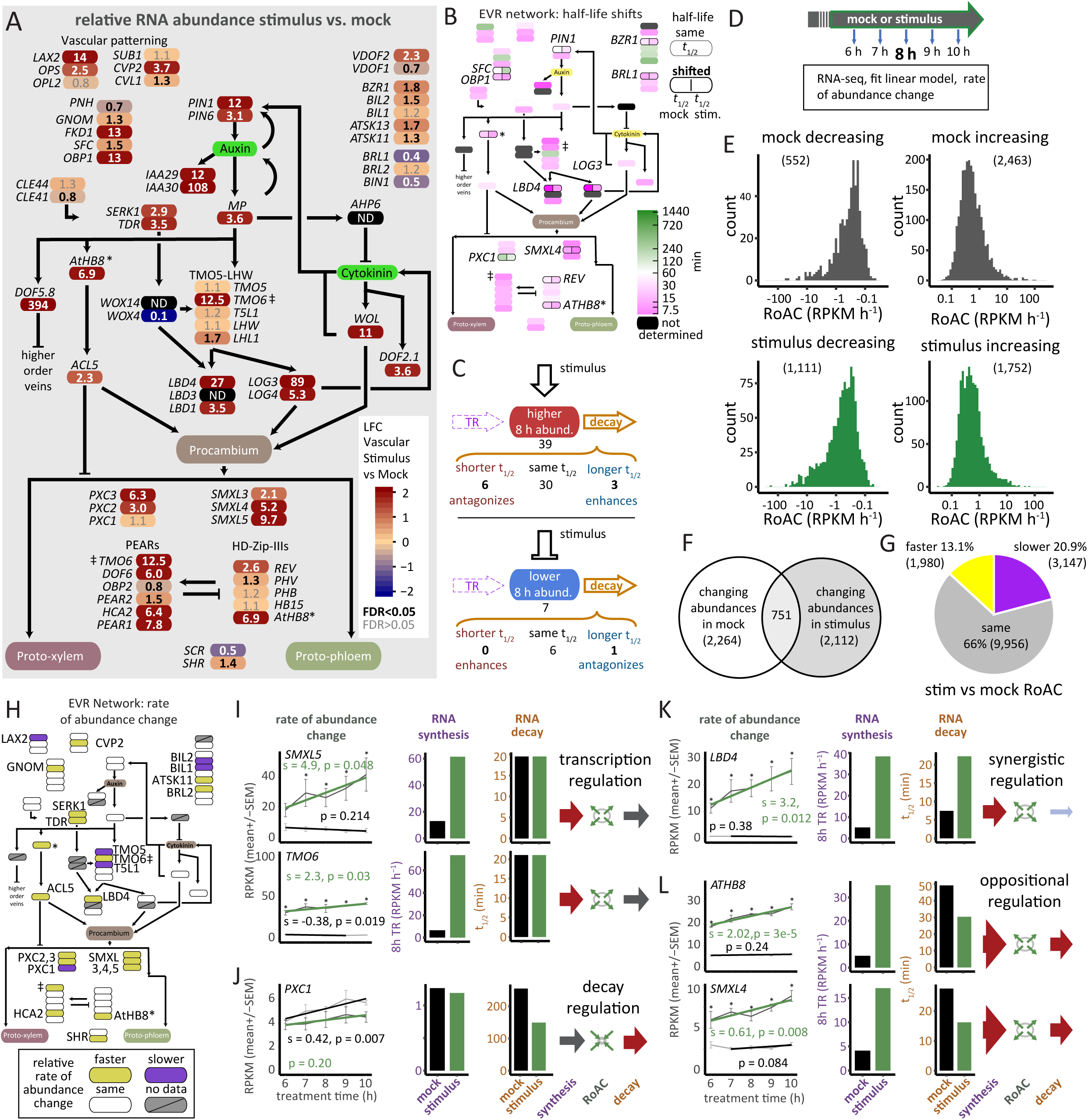
Broad regulation of RNA decay and RoAC occurs during stimulus response. (A) Effect of stimulus response on members of the early vascular regulator gene network. Heatmap of relative RNA abundance (8h stimulus vs. mock). Color and numbers indicate log2 fold change and fold change, respectively. Bold numbers indicate significant changes (FDR < 0.05). *, *ATHB8* is represented in two locations on the map. ‡, TMO6 is also represented twice. (B) *t*1/2 regulation in the EVR network (as in (A)). Color indicates *t*1/2. 11 RNAs shifted *t*1/2s and are indicated by a black outline and label. (C) Schematic of RNA regulation occurring for EVR network RNAs with higher (top) or lower (bottom) abundance (FDR < 0.05). (D) Schematic for determining rates of abundance change. Mock- and stimulus-treated tissue were collected after 6, 7, 8, 9, or 10 h to model the RoAC across the 8 h time point for each RNA. (E) Distributions of RoAC for RNAs that were decreasing (left) or increasing (right) abundance at 8 h mock (top) or stimulus (bottom). (F) Venn diagram showing RNAs with changing abundances in mock or stimulus. (G) Pie chart showing the effect of stimulus on RoAC; rates increased (faster), decreased (slower), or were the same. (H) Net effect of RNA regulatory mechanisms on RoAC (8 h stimulus vs. mock) in the EVR network (as in (A)). Positive or negative influence was found on rates of the 23 labeled RNAs. (I-L) RNAs of the EVR network reveal distinct regulatory modalities through changes in transcription and decay rates as indicated by arrows: red, faster; gray, unchanged; blue, slower.

To clarify how the 8 h *t*1/2s influenced RNA abundances, we identified RNAs with actively changing abundances across the 8 h timepoint; this used RNA abundances (RNA-seq) from 6, 7, 8, 9 and 10 h (Figure 2D) and linear regression to estimate slopes of RNA abundance change. These additional data allowed us to focus on the RNAs whose abundances were actively changing and also allowed us to calculate TR for each RNA as the amount of synthesis that was required to sustain the rate of abundance change (RoAC) occurring at 8 h, given the *t*1/2 ^20,66,67^ (Supplemental Figure S3B, S3C, S4A). These data showed that about a third of the RNAs had significant rates of abundance change (i.e., not 0, p < 0.05). These rates were broadly distributed with most RNAs changing abundance in a single condition (85%), and 15% (751) changing in both mock and stimulus (Figure 2E-2G).

We re-evaluated regulation of the 57 EVR network genes using these data; 23 of these RNAs had actively changing abundances (p<0.05)(Figure 2H). Among these 23, 19 had indistinguishable mock and stimulus *t*1/2s, indicating that transcription regulation alone was driving their changing abundances. This defined a transcriptional regulatory modality (e.g., *SMXL5* and TMO6) (Figure 2I). Another gene, *PXC1*, defined an RNA decay regulatory modality; its RNA’s much shorter stimulus *t*1/2 drove its actively changing abundance as its mock and stimulus TRs were similar (i.e., below our difference cutoff, Supplemental Figure S3B, S4B, Figure 2J). Two additional regulatory modalities featured changes in both RNA *t*1/2s and TRs. *LBD4* defined a synergistic regulatory modality in which longer *t*1/2 and faster TR both contributed to its increasing abundance (Figure 2K). *ATHB8* and *SMXL4* defined an oppositional regulatory modality. These RNAs had increasing abundances, faster TRs, and shorter *t*1/2s (Figure 2L) indicating the rate changes opposed each other. Oppositional regulation could be highly inefficient, e.g., *SMXL4*’s increasing abundance required a 1.66- fold faster TR to counter its shorter *t*1/2 (17.1 RPKM h^-1^ vs 10.3 RPKM h^1^ without the change in RNA *t*1/2). Because including RoAC data provided greater insight into the roles of RNA decay, we extended this second analytical strategy to the whole transcriptome, using the 14,600 genes with TRs, *t*1/2s and RoAC data.

### Transcriptome-wide analysis reveals RNA buffering as an additional regulatory modality

We first determined modality usage across the full transcriptome. We placed the 4,268 RNAs with actively changing abundances at 8 h into the transcription, decay, synergistic, and oppositional modalities. 3,108 had indistinguishable mock and stimulus *t*1/2s, indicating that their RNA abundances were transcriptionally regulated (Figure 3A).

**Figure 3.**
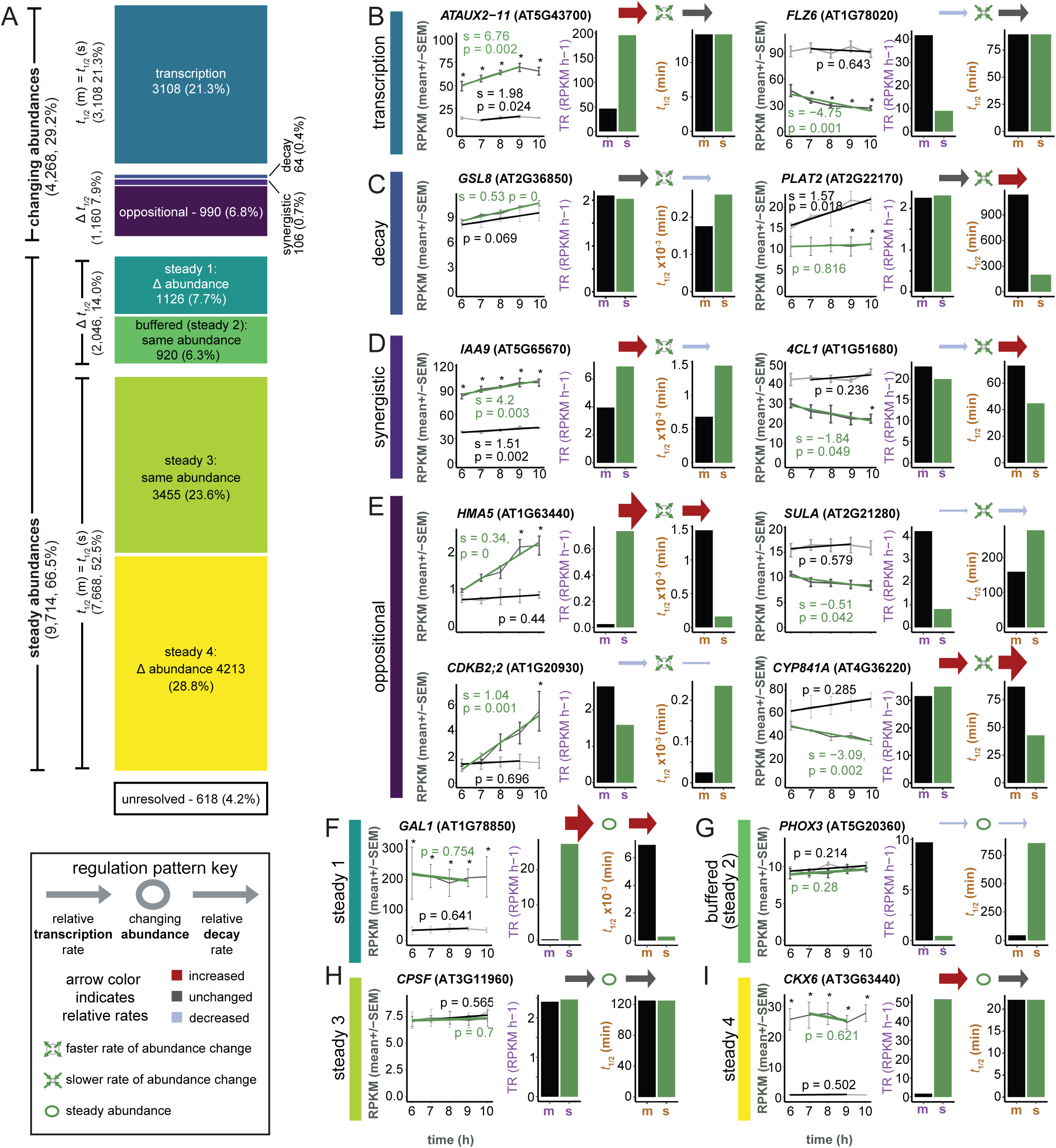
Transcriptome-wide snapshot of 8h stimulus response includes active regulatory modalities and steady abundances maintained through changes in flux. (A) Regulatory modalities are defined by change or lack of change in synthesis and decay and the net effect on RoAC. Counts (and proportions) of RNAs exhibiting behavior of each regulatory modality are presented as a stacked bar plot. (B-I) Regulation patterns of faster or slower synthesis and/or decay are demonstrated for each regulatory modality and are represented pictographically as larger red (faster TR/shorter *t*1/2) or smaller blue (slower TR/longer *t*1/2) arrows on either side of an oval. Ovals represent RNA abundance. Green arrows around the ovals represent increasing or decreasing RoAC. For each example RNA, abundance measurements (RPKM mean±SE) at 6, 7, 8, 9, 10 h of mock (thin light gray line) and stimulus (thin dark gray line) along with linear models of RoAC are shown for mock (thick black line) and stimulus (thick green line). Model parameters label the graphs: s, slope of the linear model (RPKM h-1); p, slope p- value; *, indicates a significant difference at the given time point, FDR < 0.05. *t*1/2s and TRs are also shown in bar plots to the right for mock (m), and stimulus (s).

Transcription alone was the major modality of changing abundances (21% of the full dataset) and its regulation led to both faster (1,305 RNAs) and slower (1,803) rates of abundance change (Figure 3B). Only 64 genes encoded RNAs whose abundances were regulated by decay alone (Figure 3C) making it the least used regulatory modality. The remaining 1,096 genes had RNA abundances regulated by the combined influence of transcription and decay. Synergistic regulation was found for a small number of RNAs (106) (Figure 3D) whereas oppositional regulation was far more common (990 RNAs) (Figure 3E).

We also found *t*1/2 regulation among RNAs with unchanging abundances (steady, RoAC p > 0.05). These fell into two groups based on abundance differences. One group of RNAs had differing *t*1/2s and abundances, which we called steady 1 (1126 RNAs)(Figure 3F). The other group had differing *t*1/2s but abundances were indistinguishable (steady 2, 920 RNAs) (Figure 3G and Supplemental Figure S4C). Steady 2’s combined features – changes in *t*1/2 and TR but not abundance – fit the operational definition of RNA buffering^68^. Finding RNA buffering as part of responsive gene expression was surprising because it has previously been observed in systems with dysfunction in transcription or decay machinery^34–37,41,42^.

The remaining RNAs with steady abundances and unchanged *t*1/2s also fell into two groups; steady 3 RNAs had indistinguishable abundances (3455 RNAs)(Figure 3H) and steady 4 RNAs had different mock and stimulus abundances (steady 4, 4213 RNAs)(Figure 3I).

Because steady 4’s abundances were not actively changing, the regulation leading to these abundance changes occurred prior to our experimental window. RNAs with differing abundances were maintained through different TRs (steady 4, Figure 3I) or by different RNA *t*1/2s with or without TR changes (steady 1, Figure 3F).

### *t*1/2 and TR specialization of regulatory modalities

To discover whether RNAs with short or long t1/2s were specific to certain modalities, we compared modality-specific distributions of mock *t*1/2s to that of the full transcriptome. All modalities showed similar mock *t*1/2s distributions (Figure 1L, Supplemental Figure S5A- S5E). Mock TRs also showed similar broad distributions. This indicated that none of the modalities were specialized for fast or slow rates (*t*1/2 and/or TR).

To examine whether magnitudes of *t*1/2 and TR changes differed by modality, we compared the mean *t*1/2 and TR rate changes for each (Table 1 and Supplemental Figure S5A-S5E).

**Table 1.** Magnitudes of transcription rate and half-life change by regulatory modality.

| **Modality** | **$t_{1/2}$ shift** |  | **transcription rate change** |  |
| --- | --- | --- | --- | --- |
|  | fold change mean* | fold change median** | fold change mean* | fold change median** |
| transcription | 1.0 | 1.0 | 1.8 | 1.5 |
| decay | 1.9 | 1.7 | 1.0 | 1.1 |
| synergistic | 2.3 | 1.9 | 1.6 | 1.4 |
| oppositional | 2.7 | 2.0 | 3.1 | 2.3 |
| steady 1 | 2.8 | 2.0 | 3.2 | 2.6 |
| buffered steady 2 | 2.7 | 2.0 | 2.8 | 2.0 |
| steady 3 | 1.0 | 1.0 | 1.1 | 1.1 |
| steady 4 | 1.0 | 1.0 | 1.5 | 1.4 |

\* $2^{\text{mean}(|\log_2(\text{stimulus/mock})|)}$ ; \*\* $2^{\text{median}(|\log_2(\text{stimulus/mock})|)}$

When the rate changes occurred for one process, i.e., transcription and decay modalities, the magnitudes of rate changes were very similar. This indicated that these two processes have similar capacity to impact RNA abundance. Four modalities featured changes in both transcription and decay: synergistic, oppositional, steady 1, and buffered (steady 2).

Synergistically regulated RNAs underwent small changes in *t*1/2 and TR. By contrast, oppositionally regulated, steady 1 and buffered (steady 2) RNAs underwent large changes in *t*1/2 and TR. Thus, modalities do appear to have specializations for rate changes.

### Synergistic and oppositional modalities use distinct mechanisms

We next explored gene-specific relationships between *t*1/2 and TR across the modalities. The simplest were transcription and decay; both of these modalities had rate changes that ranged from modest to large (>8-fold for *t*1/2 and >10-fold for TR) (Figure 4A-4B).

**Figure 4.**
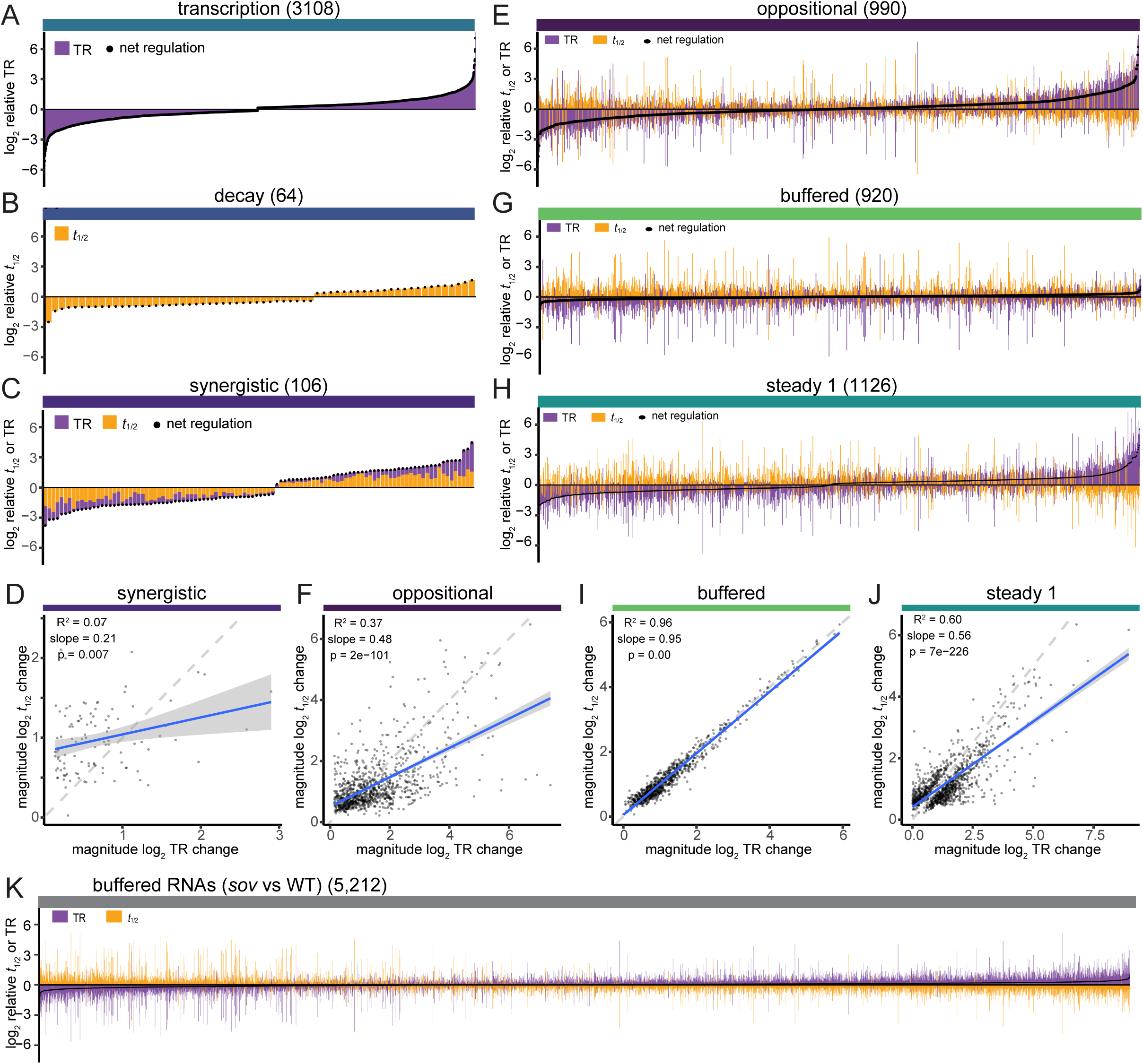
Dynamic range in transcription and *t*1/2 regulation are modality dependent. (A-G) Contributions of transcription and decay regulation for individual RNAs. Stacked bar plots show direct comparison of regulation of TR and *t*1/2 for each RNA of regulatory modalities; bars indicate direction and size of log2 relative TR (purple) and log2 relative *t*1/2 (orange). Black dots indicate net regulation (sum of the LFC rel. TR and LFC rel. *t*1/2). Genes within each regulatory modality are organized by increasing net regulation. The net regulation extremes that slightly deviate from 0 in the buffered RNAs reflects abundance variance below the cutoff.

For modalities with changes to both *t*1/2s and TRs, the gene-specific ranges of *t*1/2 and TR were represented by stacked bar plots. In the synergistic modality, the cooperativity of changes in *t*1/2 and TR was represented by bars stacked on the same side of the X axis (Figure 4C). Despite the cooperative directionality of rate changes, magnitudes of *t*1/2 and TR changes differed, suggesting that these processes were regulated independently.

Modeling of magnitudes of *t*1/2 changes as a function of magnitudes of TR changes showed low concordance; only 7% of the variance was explained by TR changes (Figure 4D).

Moreover, changes in *t*1/2 provided 65% of the total regulatory changes; that is, RNA abundance regulation of most synergistically regulated genes was driven predominantly by gene-specific change in *t*1/2s.

The oppositional modality’s stacked bar plots had a strikingly different appearance. The gene-specific ranges of *t*1/2 and TR showed high gene-to-gene variability. These changes were often very large, far exceeding that required for net regulation (the sum of log2 relative *t*1/2 and TR, indicated as black dots) (Figure 4E). In addition, for each gene, *t*1/2 and TR underwent similar rate changes (i.e., mirrored). In other words, oppositional decay was remarkable for its frequent large and opposing changes in transcription and decay that resulted in small net regulation. In modeling this modality, TR change magnitudes explained 37% of the variance of *t*1/2 change magnitude (Figure 4F). Mirroring of gene-specific rate changes was most obvious in Figure 4E’s central region where net regulation was smallest. In the upper and lower 10% of net regulation, 78% of RNAs had larger changes in TR than *t*1/2 (Supplemental Figure S5F-S5G). In other words, in oppositional regulation changing abundances can be driven by transcription or decay, but in this study, transcription was the more frequent driver.

The synergistic and oppositional modalities were similar in that both brought about changes in RNA abundances. However, they appear to do it in distinct ways. *t*1/2 regulation was independent of transcription for RNAs in the synergistic modality. By contrast, *t*1/2 regulation was linked to regulation of TR in the oppositional modality. These features suggest that synergistic and oppositional regulation are mechanistically distinct.

### Antagonistic regulation of transcription and decay is a common strategy for transcriptome homeostasis

In addition to the oppositional modality, buffered (steady 2) and steady also included opposing changes in *t*1/2s and TRs. As with oppositional, these modalities had average rate change magnitudes that were relatively large (Table 1). To assess whether buffered (steady 2) and steady 1 shared oppositional’s gene-specific rate change patterns, we analyzed gene-specific rate changes (*t*1/2 and TR) and found large ranges, with high gene-to-gene variability and commensurate changes in *t*1/2 and TR (Figures 4G-4H). Buffered RNAs are defined as those with insignificant changes in abundance despite changes in *t*1/2 and TR^68^. Accordingly, we found that TR change magnitudes explained 96% of variance in *t*1/2 change magnitudes (Figure 4I). RNAs in the steady 1 modality, which had different mock and stimulus abundances, had variance in *t*1/2 change magnitudes that were largely explained by TR change magnitudes (60%, Figure 4J). These observations, including the similar sized changes in *t*1/2 and TR for three modalities, oppositional, buffered (steady 2), and steady 1, suggest that they use a common regulatory mechanism that links regulation of decay and transcription.

To assess whether buffered RNAs in general might share these same features, we re- analyzed RNAs previously identified as buffered in the Arabidopsis *sov*/*dis3l2* mutant^40^. We examined these data using stacked bar plots and observed that magnitudes of rate changes showed high gene-to-gene variability (Figure 4K). In addition, gene-specific changes in *t*1/2 and TR had similar magnitudes. This supports buffering being part of a larger shared mechanism that includes regulation leading to altered RNA abundances.

RNAs buffered in *sov* mutants showed kinetic features suggestive of an overcompensating mechanism^39^; buffered RNAs that had shorter *t*1/2s also had slightly higher abundances and RNAs that had longer *t*1/2s had slightly lower abundances. While individually these abundance differences were insignificant, there was a strong link between the direction of abundance changes and *t*1/2 changes (Supplemental Figure S5I). To assess whether this study’s buffered RNAs (steady 2) also exhibited features of overcompensating feedback, we examined the pattern of *t*1/2 changes for RNAs with small (insignificant) abundance changes. Similar numbers of steady 2’s buffered RNAs shifted to shorter and longer half-lives in the higher and lower abundance groups indicating no overcompensating feedback (Supplemental Figure S5J). Variability in use of over-compensating feedback in RNAs buffering might reflect the how RNAs are selected for buffering; overcompensating feedback in *sov* arose because the RNA substrates of SOV shifted to degradation through decapping.

### Functional roles of regulatory modalities

Finally, we considered whether modalities might be used for specific functions, for example, during sequential waves of expression that occur during development^46,52,53^. At the beginning of a wave of expression abundances typically increase, peak, then decrease^69^. We identified RNAs in these three phases by clustering the 4,268 RNAs with changing abundances based on abundance profiles across the 6, 7, 8, 9, 10 h timepoints. We sorted the clusters into three groups: diverging abundances (1,366-genes in 11 clusters), parallel abundances (1,220 genes, 6 clusters) and converging abundances (1,682-genes, 7 clusters)(Figure 5A). We found enrichment of synergistically and oppositionally regulated RNAs in the diverging clusters, and strong depletion of oppositionally regulated RNAs in the converging clusters (Figure 5B). This suggests roles for both synergistic and oppositional modalities in driving diverging abundances.

**Figure 5.**
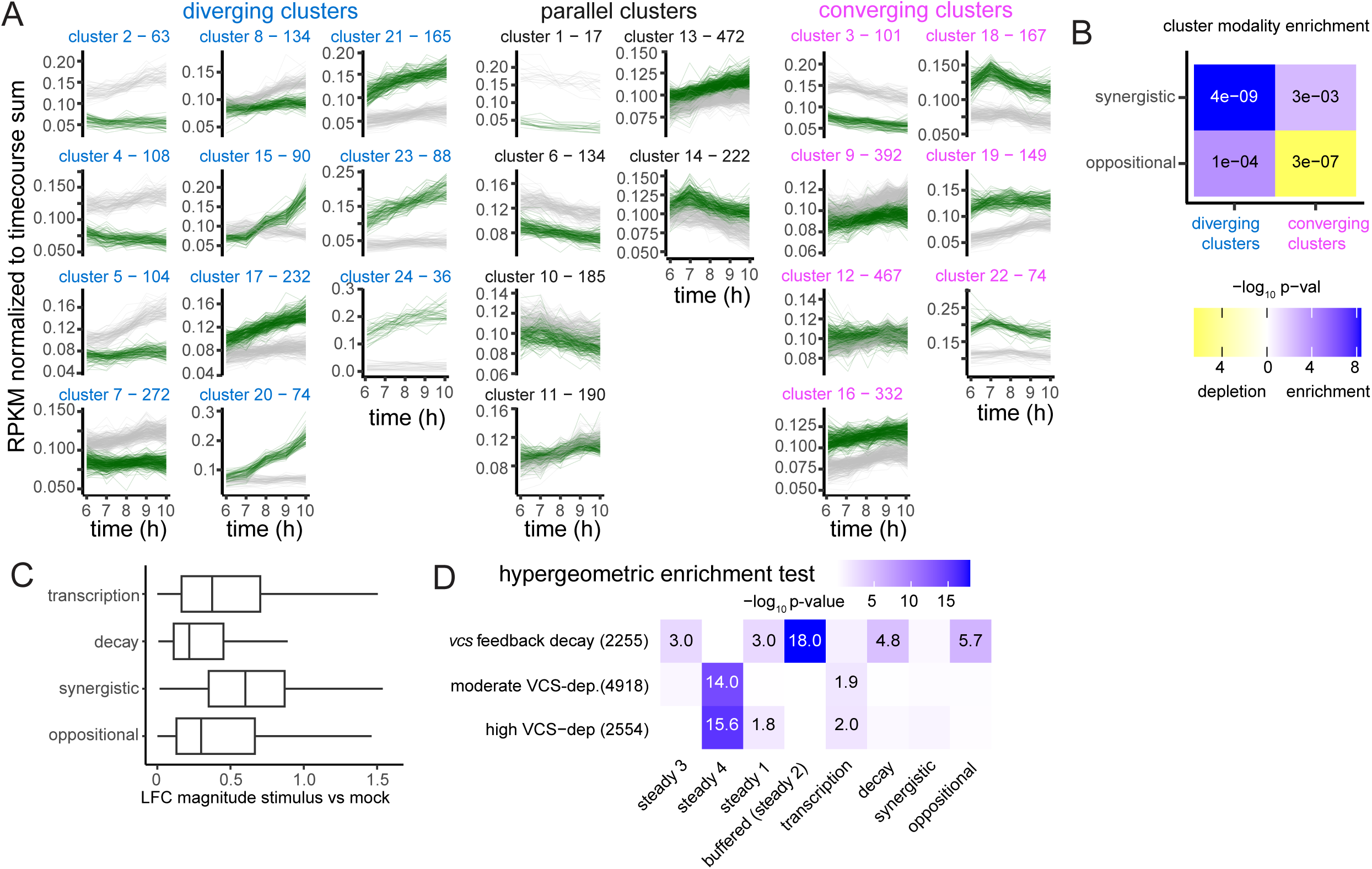
Synergistic and oppositional modalities drive responsive RNA abundance. (A) RNA abundance response profiles. k-means clustering of 4,268 RNAs of the transcription, decay, synergistic, and oppositional modalities. Abundance data of each RNA was normalized to the total (sum) abundance of all time points so clustering would group RNAs with similar abundance response profiles independent of level of expression. Clusters were numbered based on the relative abundance difference between mock (gray lines) and stimulus (green lines). Cluster number and size label each plot; label color indicates abundance trajectories at 8 h: diverging (blue), converging (pink), or other (black). (B) Modality enrichment among responsive abundance. RNAs of diverging or converging abundance clusters in (A) were grouped and evaluated for enrichment (blue) or depletion (yellow) of the synergistic and oppositional regulatory modalities by the hypergeometric test; enrichment/depletion are displayed as a heatmap of and labeled with -log10 p-values. (C) Box and whisker plot of the abundance change magnitudes (|L2FC abundance|) by regulatory modality. Abundances of all stimulus time points were compared to those of all the mock time points. (D) Enrichment analysis of RNA groups with distinct changes in *t*1/2 in *vcs* mutants: *vcs* feedback decay (rel. *t*1/2 *vcs*/WT < 1), moderate *VCS*−dependence (rel. *t*1/2 *vcs*/WT > 1 & < 2), high *VCS*−dependence (rel. *t*1/2 *vcs*/WT > 2). Significant enrichments are labeled with -log10 p-values.

If these modalities are important for diverging abundances, RNAs in the synergistic and oppositional modalities might show substantial changes in abundance. Although this analysis was complicated by abundances that were in the process of changing, on average RNAs in the synergistic modality showed the largest abundance changes (Figure 5C).

Together, these data support the synergistic modality as being the most important effector for large changes in RNA abundance. Furthermore, because abundances of synergistically-regulated RNAs were primarily driven by decay, RNA decay emerges as an important regulator of RNA abundances.

We also assessed potential specialized functions of the modalities using GO enrichments^70^. The strongest of these were for RNAs in steady 4, which had differing mock and stimulus abundances that arose prior to the experimental (Supplemental Figure S5K). Steady 4 RNAs were enriched in a set of GOs related to ribosome constituents and translation, suggesting that ribosomal protein RNA abundances were adjusted early and without contributions by decay regulation.

RNA decay pathways have specialized functions^40,60,71^, and we considered that decay pathways might also have specialized contributions to modality regulation. To test this idea, we used *t*1/2 dependence on VCS/EDC4 (the proportion of decay that was lost in *vcs*/*edc4* mutants, which lack the mRNA decapping complex scaffold^40^). We compared enrichment of three sets of genes: sets with high dependence (stabilized > 2-fold in *vcs* mutants), moderate dependence (stabilized < 2-fold in *vcs* mutants), and a set which were strongly destabilized in *vcs* mutants which we referred to as feedback regulated^40^. The RNAs in steady 4 were enriched in both the high and moderate VCS-dependent RNAs, indicating an important role for mRNA decapping in prior abundance regulation or in facilitating maintenance of steady RNA abundances (Figure 5D). The strongest enrichment was for buffered (steady 2) RNAs; these were enriched in *vcs* feedback-regulated RNAs. Thus, this group includes RNAs that appear to have a propensity for feedback regulation.

## DISCUSSION

Measurable changes in mRNA abundances, commonly referred to as gene expression, are the outcomes of changes in rates of transcription and RNA decay. Here, we demonstrate that RNA decay played important roles in the regulation of RNA abundances in response to a developmental stimulus. First, we demonstrated that decay and transcription had similar capacities to adjust RNA abundances. The decay capacity was showcased by the regulation of RNAs in the synergistic modality; these RNAs showed gene-specific stimulus-induced changes in *t*1/2 and TR that were cooperative, led to the largest abundance changes, and were primarily driven by decay. Oppositionally regulated RNAs showed stimulus-induced changes in *t*1/2 and TR that were antagonistic, they resulted in modest abundance changes, and while transcription was the main driver, decay-driven RNA abundance changes were also observed. One function of oppositionally-regulated decay was to temper excessive changes in TRs and thus facilitated transcriptome homeostasis. In oppositionally regulated RNAs, *t*1/2 and TR changes had similar or mirrored magnitudes in contrast to synergistically regulated RNAs which showed *t*1/2s that were regulated independently of TRs. Because of this and the difference in average modality rate changes, the synergistic and oppositional modalities appear to be governed by distinct mechanisms. Moreover, buffered (steady 2) and steady 1 RNAs had large and mirrored rate change magnitudes that were similar to oppositionally regulated RNAs. This suggested that these three groups had a shared regulatory mechanism that accounted for 90% of the RNAs with regulated *t*1/2s; we call this shared mechanism countercyclical regulation.

Synergistic and oppositional relationships between rates of transcription and decay appear to have specialized usage. In yeast, RNAs exhibiting behavior similar to these modalities showed stress-specific roles; response to DNA damage included strong use of synergistic-like regulation while responses to osmotic and oxidative stress predominantly used oppositional-like regulation^18,72,20^. Similarly, studies using drosophila and mouse cells found widespread use of oppositional regulation while synergistic appeared to have more specialized functions, e.g., in cell-type specific regulation^18,20,73,74^. Here, the synergistic modality was used for large changes in expression, including in genes whose RNAs were becoming differentially regulated. Consistent with the idea that modalities are used during specific responses, oppositional regulation is used as RNA abundances change in response to specific stimuli in yeast and animals cells^20,75^. In our study, synergistic and oppositional regulation also played roles in distinct phases of gene expression, such as the sequential waves of gene expression associated with vascular transdifferentiation^76,51,77,45,52,53^. RNAs with diverging abundances were more enriched in synergistically and oppositionally regulated RNAs as compared to RNAs with converging abundances. Whether other sorts of signaling change the balance between use of synergistic and oppositional regulation awaits further study.

Countercyclical regulation of RNAs was apparent in the oppositional, buffered (steady 2) and steady 1 regulatory modalities. We initially focused on RNA abundance change and placed oppositional and buffered RNAs in distinct groups. However, by prioritizing their regulation, RNAs of these modalities could be arrayed across a continuum, with the changing abundances of oppositionally-regulated RNAs defining the ends of the continuum and buffered RNAs occupying the center. Steady 1 RNAs would also reside at the ends of the continuum but differ from oppositionally-regulated RNAs because their different abundances have reached steady state. This suggests that their different abundances might have arisen from a countercyclical mechanism, and that these rate changes were durable. In contrast to the global nature of RNA buffering in yeast^79^, countercyclical regulation appeared to be gene specific; magnitudes of *t*1/2 and TR changes differed between regulated genes and only a portion of the transcriptome was regulated in this manner. Gene-specific buffering was observed in studies of mouse embryonic stem cells^78^ and also occurred in *sov* mutants of Arabidopsis, where 42% of the transcriptome was affected^40^.

A future challenge will be to determine the mechanism of RNA buffering. One model for RNA buffering features nuclear-cytoplasmic shuttling of RNA buffering effectors. This model is supported by studies in yeast, where XRN1 shuttling plays an important role^34,36–38,80^. Cytoplasmic XRN1 degrades decapped and fragmented mRNAs^37^ and shuttles into the nucleus in a decay-dependent manner. Nuclear XRN1 binds DNA and contributes to gene expression^80^. This model is consistent with the global RNA buffering observed in yeast, however plants lack a cognate homolog of XRN1 and instead use the XRN4 paralog which lacks key nuclear localization signals^62,80,81^. Other effectors might be conserved in plants such as those that shuttle signals dependent on 3’ to 5’ decay^42^ but how this model explains gene-specific buffing is unclear. A second model comes from the observed mirrored changes in *t*1/2 and TRs which suggest that coordinated *t*1/2 and TR regulation.

Potential mechanism for coordinated regulation include mRNA imprinting (e.g., with Rpb4 and Rpb7)^33,72,82–84^, gene-specific mRNA methylation linked to transcription elongation rates^74^, and a releasing-shuttling mechanism that coordinates regulation^85^. Finally, it is also possible that the mirrored *t*1/2s and TRs are simply the outcomes of default rapid RNA degradation in the absence of stabilization. A rapid increase in transcription might produce RNAs in excess of the stabilization capacity (e.g., from a stabilizing RNA binding protein), potentially leading to similar changes in *t*1/2s and TRs. For example, an RNA binding protein that stabilizes RNAs might become saturated after a transcription response, leading to excess unbound RNAs being targeted for degradation^86^. This scenario is consistent with transcription being the primary driver of RNA abundance change in oppositional decay. If correct, then abundance regulation of stabilizing RNA binding proteins might explain RNA buffering’s maintenance of normal RNA abundances.

A surprising outcome of this study was finding buffered RNAs among the transcriptome responses to a developmental stimulus. RNA buffering is widely described as a response to dysfunction in RNA decay and transcription, whether caused by mutation or pharmaceuticals^34,35,74^, and seldom sought in studies of development. However, an implication of this finding is that a developmental stimulus can evoke transcriptome responses that are interpreted as abnormal and thereby evoke RNA buffering. The need for measurements of RNA decay rates to diagnose RNA buffering likely means that RNA buffering might continue to be a cryptic feature of many studies.

Countercyclical regulation is a mechanism for transcriptome homeostasis because it reins in RNAs whose *t*1/2s and/or TRs are excessive. Here we found RNA buffering as part of a stimulus response, suggesting that the stimulus response included excessive *t*1/2 and/or TR responses. In yeast, RNA buffering is the key homeostatic mechanism; it maintains stereotypical mRNA abundances when components of the transcription or decay machinery are dysfunctional^34,35,87–90^. Yeast also show oppositional regulation, however whether it also uses a countercyclical mechanism is unknown. Countercyclical-like regulation has also been documented in drosophila and mammalian cells^34,35,73,74,87–90^ and so it appears to be an evolutionarily conserved mechanism that places the regulation of decay at the center of both RNA abundance regulation and transcriptome homeostasis.

## Acknowledgements

We thank the Arabidopsis Biological Resource Center for maintaining and sharing genetic material. We thank Dominique Gagliardi and members of the RNA degradation group at the Institut de Biologie Moléculaire des Plantes (IBMP) in Strasbourg, France and Sally Assman from the Pennsylvania State University Huck Institute of the Life Sciences for critical suggestions. This work was supported by the National Science Foundation under award numbers 1838345 and 1616779. R.S.S. was supported by an NIH Developmental Biology Training Grant (5T32 HD07491). Research reported here used the High-Throughput Genomics and Cancer Bioinformatics Shared Resource at Huntsman Cancer Institute at the University of Utah, supported by the National Cancer Institute of the National Institutes of Health under award number P30CA042014. The content is solely the responsibility of the authors and does not necessarily represent the official views of the NIH.

## Data availability

Sequencing data for this study can be accessed at the Gene Expression Omnibus (https://www.ncbi.nlm.nih.gov/geo/) under series accession number GSE278544.

## Author contributions

Conceptualization, Funding Acquisition, Methodology, Writing, R.S.S. and L.E.S.; Investigation, Formal Analysis, Data Curation, Visualization, R.S.S.; Supervision, L.E.S.

## Limitations of the study

RNA *t*1/2 estimates could not distinguish cytoplasmic RNA decay from decay that occurs in the nucleus and so the extent to which symmetric changes in transcription and decay rates reflected coordination between cytoplasmic and nuclear compartments is unknown. *t*1/2 measurements used leaf tissue, which resulted in the averaging over multiple cell types. Rates of abundance change were based on best linear fit across three to five timepoints, and a non-parametric fit might have yielded different estimates with potential impacts extending to TR calculations. While this study identifies transcriptome homeostasis as an important role of RNA decay, its potential link to scaling of mRNA abundance to cell size was not considered. Finally, this study’s conclusions are based on analysis of a single timepoint of a response to a single stimulus which may limit the generality of our conclusions.

## METHODS

### Plant Genetic Material and Growth Conditions

*AtHB8::4xYFP* (Columbia-0 genetic background) ^49^ were received from Arabidopsis Biological Resource Center (CS2106110, Ohio State University). Seeds were sown on moistened soil (Sungro Redi-Earth Plug and Seedling propagation mix) and imbibed in the dark at 4°C. After 2-4 d, pots were grown under continuous illumination with 75 µmol photons m^-2^ s^-1^ at 21°C for 10 d. Leaves 1 and 2 were used in all experiments unless otherwise indicated.

### Phloroglucinol Staining & Microscopy

Lignin staining was performed after clearing leaves in warm 70% ethanol. Cleared tissue was stained and imaged in phloroglucinol stain (prepared as a fresh 2:1 mixture of 3% w/v phloroglucinol in ethanol with concentrated hydrochloric acid (37 N))^91^. For DIC imaging, treated leaves were cleared using 30% acetic acid 70% ethanol overnight to 7 d, rinsed 3 times in 70% ethanol and transferred to ClearSee solution (10% w/v xylitol, 15% w/v sodium deoxycholate, 25% w/v urea)^92^, incubated overnight and transferred to ClearSee with 12.5% glycerol for mounting and imaging. Microscopy was performed on an Olympus BX-50 compound microscope. DIC images were taken with the Olympus DP-74 camera. YFP fluorescent images were captured from leaves mounted in water and using the U-MWIB filter cube by the Olympus DP-70 camera.

### Developmental Stimulus Treatment and RNA Decay Analysis

For the developmental stimulus treatment, we used the protocol of Kondo et al.^44^, and modified it for treatment to whole leaves. Detached leaves (∼18-20/sample) were floated on distilled water during collection, followed by incubation in 10% bleach 0.1% Silwet L-77 for 3-5 min, and rinsing 3 times in sterile distilled water to surface sterilize. Sterile leaves were transferred to incubation buffer (0.22% Murashigie and Skoog salts (Caisson Laboratories) [pH 5.7], 5.0% glucose, 0.2% dimethyl sulfoxide) with (stimulus) or without (mock) 20 µM bikinin (Calbiochem), 2.5 µg mL^-1^ 2,4-Dichlorophenoxyacetic acid (Sigma), and 0.5 µg mL^-1^ kinetin (Sigma). Samples were incubated at 21°C with shaking (120 RPM) for designated times and collected. To inhibit transcription, leaves were incubated in the same buffer with 1 mM cordycepin (Chengdu Biopurify Phytochemicals) and incubated with shaking for the additional designated time. Time points were selected to be equidistant on a log scale that included a short time point (15 min) for higher accuracy of short *t*1/2s. Following incubation, excess moisture was removed, and tissues were flash frozen, ground to powder and stored at -80°C until RNA isolation.

### RNA Sample processing

RNA was extracted using the Quick-RNA Plant mini-prep kit (Zymo) and Treated with DNase I. For quantitative polymerase chain reaction measurement of specific mRNA abundances, RNA was primed with random hexamers and reverse transcribed using the High Capacity cDNA Reverse Transcription (Applied Biosystems) kit. qPCR was performed on a C1000 Touch Thermal Cycler with the CFX-96 Real-Time Detection System (Biorad) using primers UBC10_qF: tcactggcaggcaacgataatgg, UBC10_qR: aagatgtcgaggcaaatgctaccg, DOF5.8_qF:cggtggtgtttcccgtaaaagc, DOF5.8_qR:cgtggcgggtatttgagcaatg, AtHB8_qF:caagggtcgcggagatcctaaag, AtHB8_qR:tctctgccctaacgaaatgaggaga, SMXL5_qF:taagccttcatgcgacaagtggac, SMXL5_qR:agaacgccccaaatcaagagtgac.

For RNA sequencing, total RNA was submitted to the Huntsman Cancer Institute High Throughput Genomic Core for library construction and sequencing. rRNAs were depleted from the samples using Ribo-Zero Plant. Stranded RNA sequencing libraries were prepared using the TruSeq Stranded Total RNA Kit (Illumina). For the RNA decay samples, purified libraries were evaluated using an Agilent Technologies ScreenTape assay. The molarity of adapter-modified molecules was defined by quantitative PCR using the Kapa Library Quant Kit (Kapa Biosystems). Individual libraries were normalized to 10 nM and equal volumes were pooled in preparation for Illumina sequence analysis. Sequencing libraries (25 pM) were chemically denatured and, for the RNA decay experiment samples,, applied to an Illumina HiSeq v4 single read flow cell. Hybridized molecules were clonally amplified and annealed to sequencing primers with reagents from an Illumina HiSeq SR Cluster Kit v4- cBot. Following transfer of the flowcell of an Illumina HiSeq 2500 instrument, a 50 cycle single-read sequence run was performed using HiSeq SBS Kit v4 sequencing reagents. For the time course experiment, libraries were transferred to the flowcell of an Illumina NovaSeq 6000 instrument, a 151 x 151 cycle paired end sequence run was performed using a NovaSeq 6000 S4 reagent Kit v1.5 (20028312).

### Bioinformatic Analysis

For analysis of RNA sequencing data, steps were performed within R using the *systemPipeR* package^93^. 50 bp reads were trimmed of adaptor sequences (Read 1 Adaptor: AGATCGGAAGAGCACACGTCTGAACTCCAGTCA; Read 2 Adaptor: AGATCGGAAGAGCGTCGTGTAGGGAAAGAGTGT) using the *trimLRPatterns* function from the *Biostrings* package^94^, and aligned to the TAIR10 genome sequence using *Tophat Bowtie2* allowing a single base mismatch^95,96^. Read counting was performed using the *summarizeOverlaps* function of the *GenomicAlignments* package^97^ using gene models of Araport11_GFF3_genes_transposons.201606.gff (accessed at TAIR: www.arabidopsis.org/download). Differential Expression analysis was performed using the *edgeR* package^98^, significant RNA abundance differences were filtered based on an FDR < 0.05. Gene alias (gene_aliases_20201231.txt) and functional descriptions (Araport11_functional_descriptions_20201231.txt), and gene ontology assignment (ATH_GO_GOSLIM.txt) files were also downloaded from TAIR. The gene ontology annotations file (go-basic.obo) was accessed at geneontology.org. Gene ontology enrichment was calculated using the *topGO* package^99^. Visualization of data used the *ggplot2* package^100^.

### RNA Kinetic Analyses

<u>RNA decay rates:</u> RNA decay rates were estimated by modeling the measured RNA abundance decrease following treatment with cordycepin to inhibit transcription^40^. RNA counts were first normalized to the total number of read counts in the library and scaled to 1 million. Scaled read counts were further scaled to a normalized gene length of 1000 nt (RPKM values). Prior to modeling, low abundance RNAs were removed (sum of the mean T0 abundances (RPKMs) of mock and stimulus were less than 2 were removed. The read counts from libraries from RNA from samples collected after cordycepin treatment were also scaled so that the mean RNA abundance of 30 stable RNAs (ATCG00490, ATCG00680, ATMG00280, ATCG00580, ATCG00140, AT4G38970, AT2G07671, ATCG00570, ATMG00730, AT2G07727, AT2G07687, ATMG00160, AT3G11630, ATMG00060, ATCG00600, ATMG00220, ATMG01170, ATMG00410, AT1G78900, AT3G55440, ATMG01320, AT2G21170, AT5G08670, AT5G53300, ATMG00070, AT1G26630, AT5G48300, AT2G33040, AT5G08690, AT1G57720) was the same. Decay rates were modeled from normalized data using the *RNAdecay* package^40,101^. We interpreted the initial decay rate estimate as representative of cellular RNA stability (in fraction of RNAs degraded min^-1^), and these were converted to *t*1/2s (min ^-1^). For identification of shifted RNA decay rates between mock and stimulus, we compared distinct models in which treatment parameters were constrained or in which they could vary independently. These models were compared using the corrected Akaike Information Criterion (AICc) metric. The model with the lowest AICc was selected and parameter estimates were used for downstream analysis.

Following modeling, RNAs with high modeled variance were filtered (σ^2^ > 0.0625).

<u>Rates of RNA Abundance Change</u>: The rate of RNA abundance change was modeled from RNA abundances measured by RNA-seq from samples treated in mock and stimulus conditions for 6, 7, 8, 9, or 10 h. Linear regression was applied using RPKM values of the 7, 8, 9 h; 6, 7, 8, 9 h; 6, 7, 8, 9, 10 h; or 7, 8, 9, 10 h. The model with the smallest stimulus slope p-value was used for downstream analysis. Modeled slopes of RNAs with slope p-values < 0.05 were considered significant and were interpreted as the rate of RNA abundance change in units of RPKMs h^-1^. Abundances of other RNAs (p > 0.05) were considered unchanging.

Transcription Rates: The TR at 8 h treatment was calculated using the measured abundance at that time point (*m*), modeled decay rate (*α*), and rate of abundance change 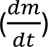 using the following equation:

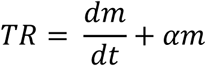

(described in Pérez-Ortín et al.^66^ and used by Xu et al.^67^).

To identify RNAs that were regulated predominantly through decay regulation, we required mock and stimulus TRs to be very close by applying a stringent filter (|log2 relative TR stimulus versus mock| < 0.138), which was selected based on the distribution of log2 relative TR (stimulus vs mock) (Supplemental Figure S4B). This also resulted in consideration of relative TR differences that were small in other modalities. 206 RNAs with calculated TRs that were negative (indicating a faster decrease in abundance in the time course experiment than could be accounted for by the measured *t*1/2) were removed. For calculating sov TR, we did not have rate of abundance change data, so we calculated the TR with the assumption that all RNAs were at a steady state (i.e, dm/dt = 0).

For clustering time course abundance profiles, abundances were normalized to the sum of the all time points in the mock and stimulus for each gene. RNAs were then clustered using the normalized values using k-means (the kmeans function from the *stats* package) clustering in R. All other statistical tests were performed using R functions.

### Identification of cell-type cluster markers from whole leaf single-cell RNA-seq

We used the dataset of whole leaf single cell transcriptomes generated Kim et al.^58^ to characterize the RNA abundance differences observed at 8 h stimulus treatment. Kim et al. ^58^ characterized 19 cell clusters among 5,230 single cell transcriptomes and associated them with distinct cell-type identities. We performed UMAP dimensional reduction and clustering analysis on their dataset using the *Seurat* package version 4.0.5^102^ to reconstitute their clusters and identify cluster markers (accessed Aug. 1, 2021 from https://www.ncbi.nlm.nih.gov/geo/: accession number GSE161332; files: GSE161332_features.tsv, GSE161332_barcodes.tsv, leaf_cell_info.txt, GSE161332_matrix.mtx). Cluster gene membership after reanalysis was not identical to those of Kim et al.^58^ but yielded a similar UMAP plot (Figure S1G) and 18 clusters which we numbered based on membership similarity with Kim et al.^58^. In our analysis, cluster 7 and 12 from Kim et al.^58^ merged, but on average clusters shared 95% membership (Supplemental Data File 3). We used the Seurat package *FindAllMarkers* function to apply the Wilcoxon Rank Sum test to RNAs present in at least 25% of cells, and with |log2FC| > 0.25 in the cell cluster compared with all others. We identified 7,996 RNAs as positive (enriched) or negative (depleted) cluster markers (Supplemental Data File 4).

---

**Supplemental Table 1.**
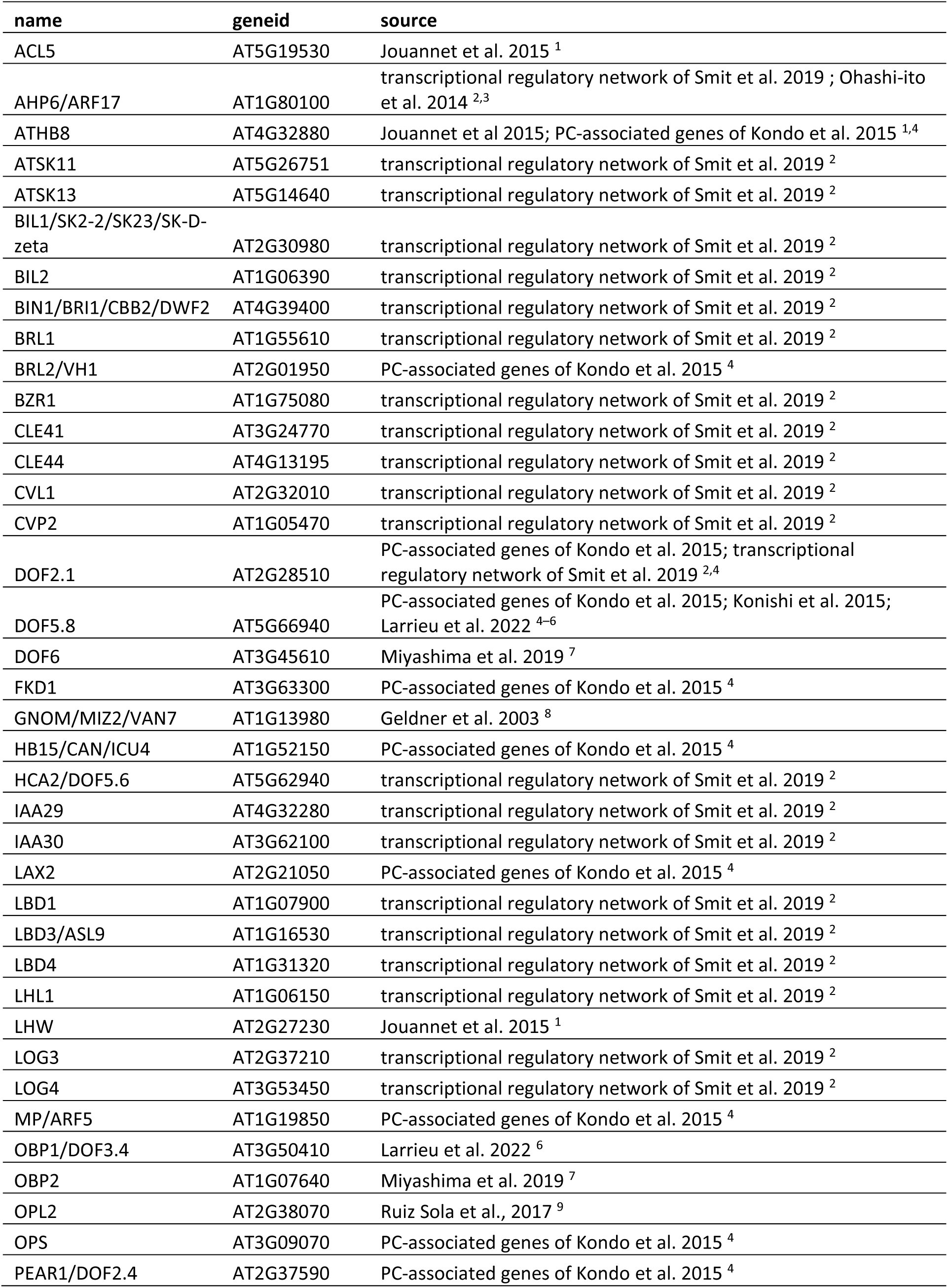
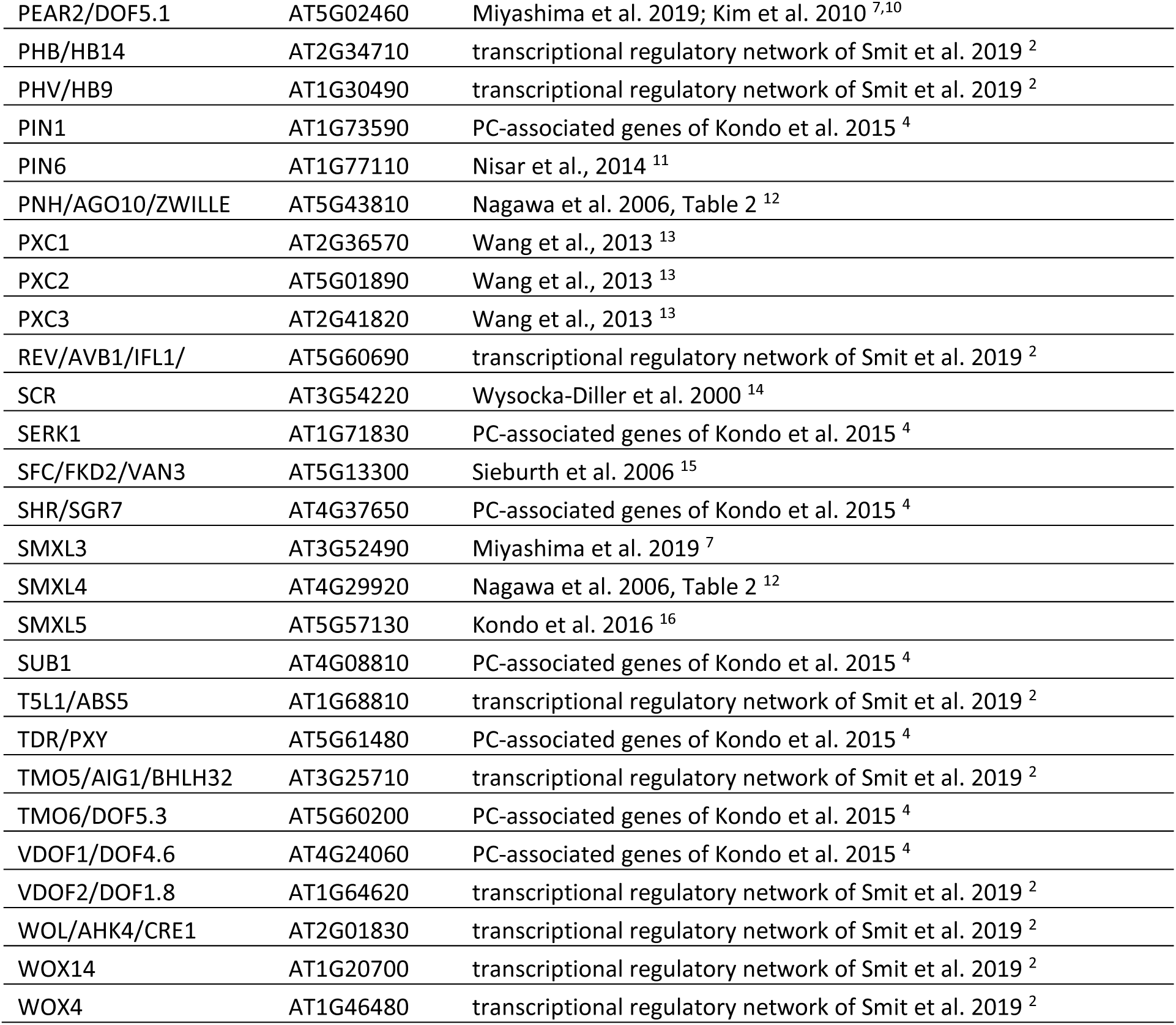
Early vascular network members and source literature.

**Supplemental Figure S1.**
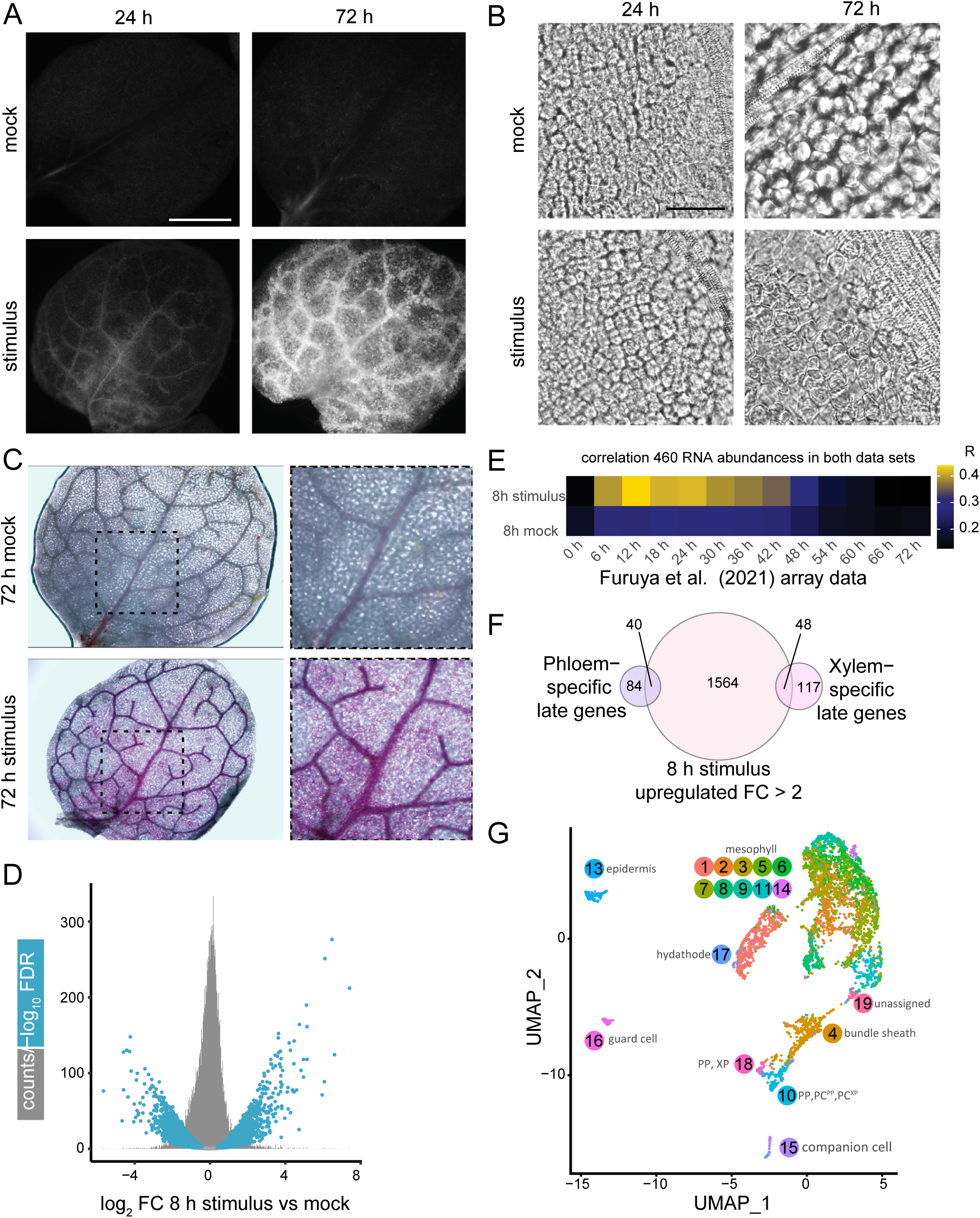
Stimulus induces developmental, molecular and biochemical changes to cells. (A) YFP fluorescence of rosette leaves 1 and 2 of *proATHB8::4xYFP* plants incubated in mock and stimulus treatments for 24 and 72 h (bar = 1 mm). (B) Development of xylem element characteristics in cells outside of vascular bundles after 72 h of stimulus treatment but not 72 h mock treatment. DIC micrograph of ectopic formation of spiral patterned secondary cell walls^103^ can be seen (bar = 50 µm). (C) Lignification of cell walls 72 h after stimulus treatment. Phloroglucinol was used to stain lignin (pink color). Box (dashed line) in the left column are enlarged to the right. (D) Volcano plot overlayed on a histogram of 8 h RNA-seq comparison of stimulus vs mock. (E) Heatmap of Pearson correlation values comparing similarity of abundance change of 460 RNA detected in both RNA seq data of this study and those measured by microarray by Furuya et al.^46^. (F) Venn diagram comparing RNA identities of 8 h stimulus-upregulated RNAs from this study and stimulus-upregulated xylem- and phloem-specific RNAs after 72 h^45^. (G) Leaf single cell RNA-seq UMAP clusters based on RNA expression measured by Kim et al.^58^. PC, procambium; PP, Phloem parenchyma; XP, xylem parenchyma. Compare Figure 1E, 1F.

**Supplemental Figure S2.**
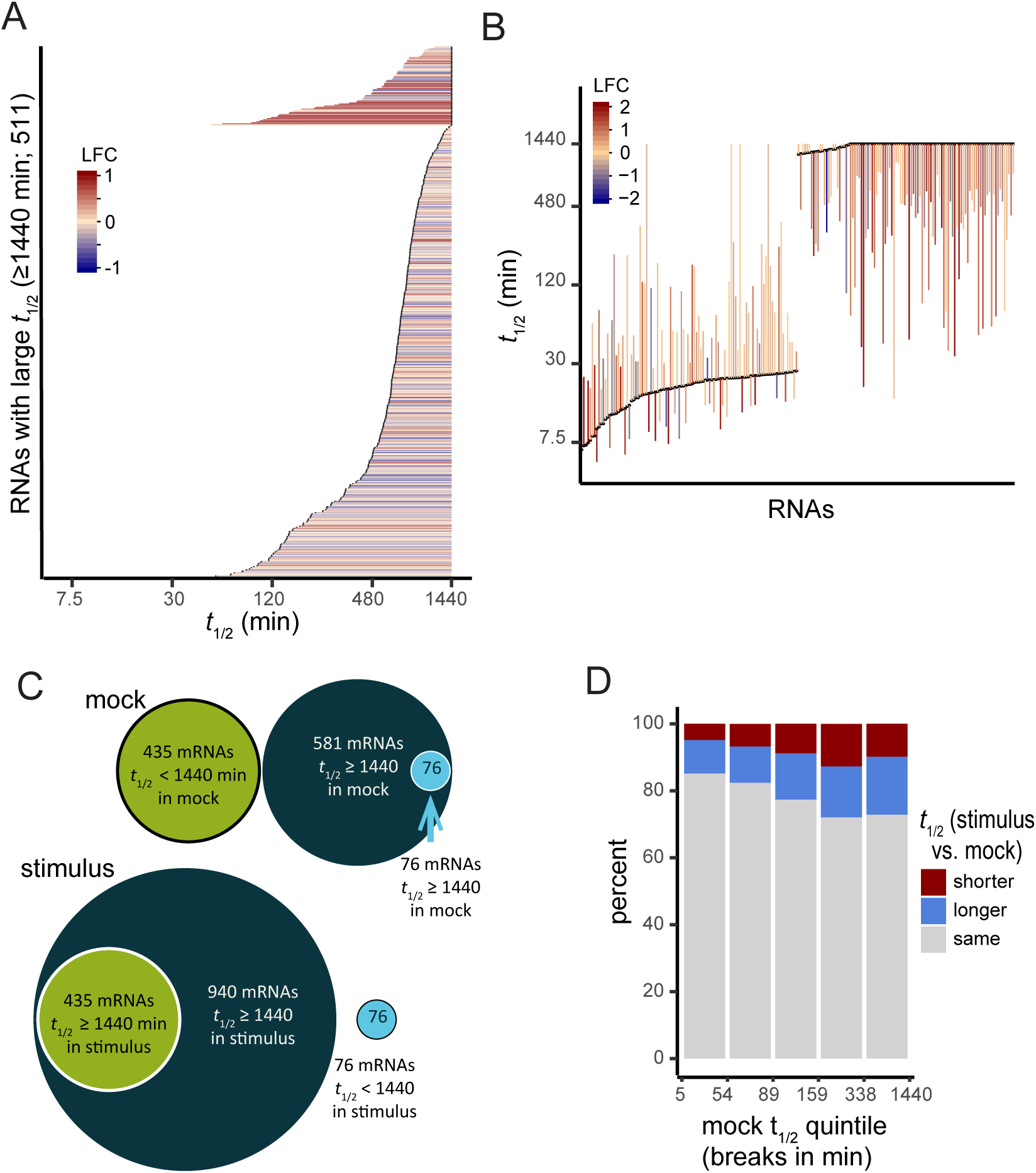
RNA t1/2 extremes have unique features. (A) Dynamic ranges of RNAs that shift to or from extremely long *t*1/2s (>1440 min) were separated from other RNAs. Line segment ends indicate *t*1/2s; black dots indicate the mock ends. (B) At *t*1/2 extremes, short short-lived RNAs tended to become longer-lived, and long-lived RNAs tended to become shorter-lived. The 100 shortest and 100 longest *t*1/2s in mock are shown. Line segments represent shifts in *t*1/2 and are positioned at the mock *t*1/2s (black dots). Line color indicates log2 relative abundance (LFC); compare with (B) and Figure 4A. (C) Dynamics of long-lived RNAs (*t*1/2 ≥ 1440 min). Some RNAs are long-lived in mock but move out of this group in stimulus (blue circle), others become long-lived in stimulus (green circle), and others remain at *t*1/2 ≥ 1440 min in mock and stimulus. (D) Bar plot comparing proportion of mock *t*1/2 quintiles with shifts in decay.

**Supplemental Figure S3.**
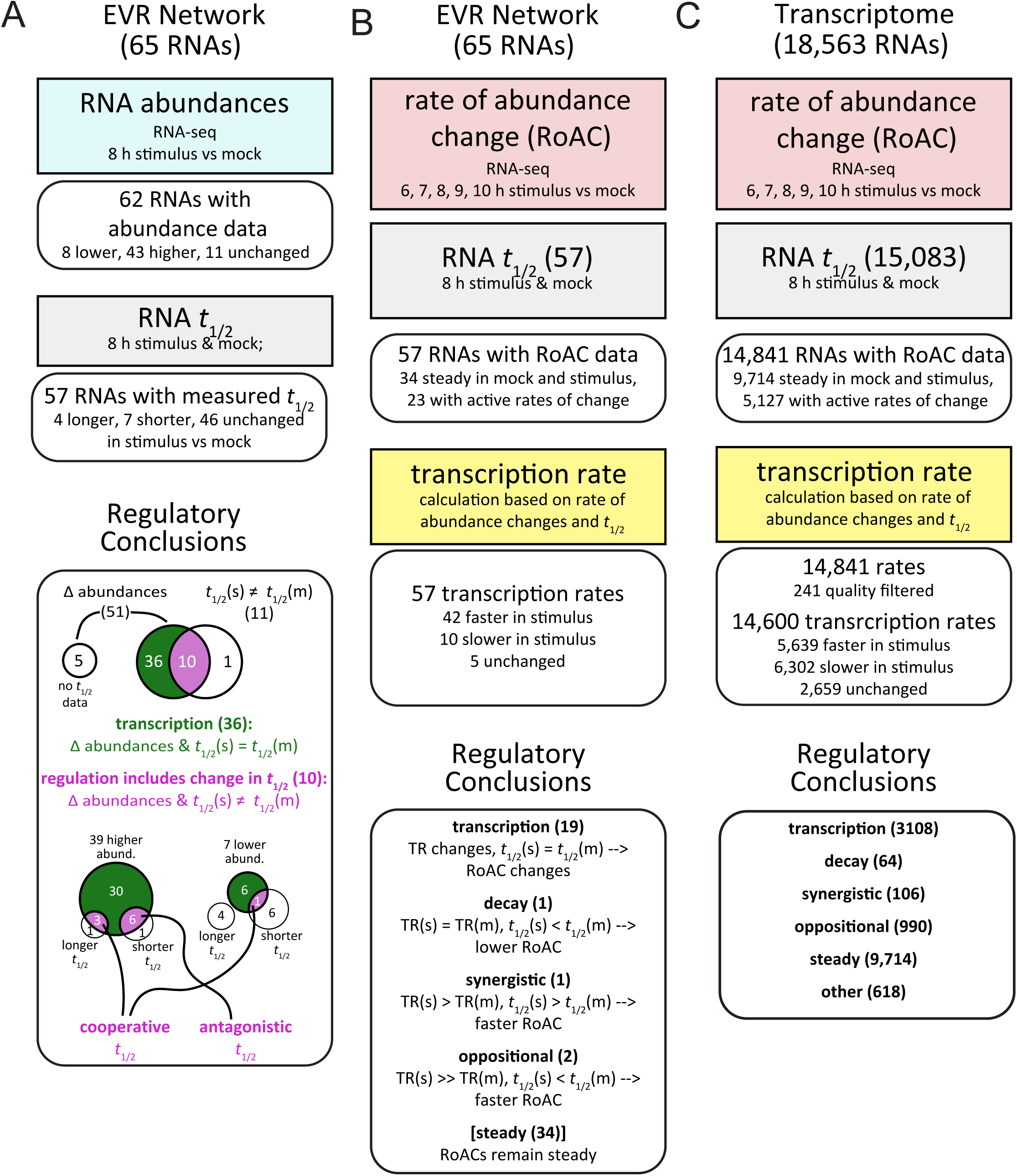
Analysis flowchart for comparing abundances, *t*1/2s, and TRs of mock and stimulus treatments. (A) Initial EVR network analysis by comparison of 8 h abundances with *t*1/2s. (B) EVR network analysis using RoAC, *t*1/2 regulation, and calculated TRs. (C) Whole transcriptome regulation analysis as in (B). See (B) for modality assignment criteria.

**Supplemental Figure S4.**
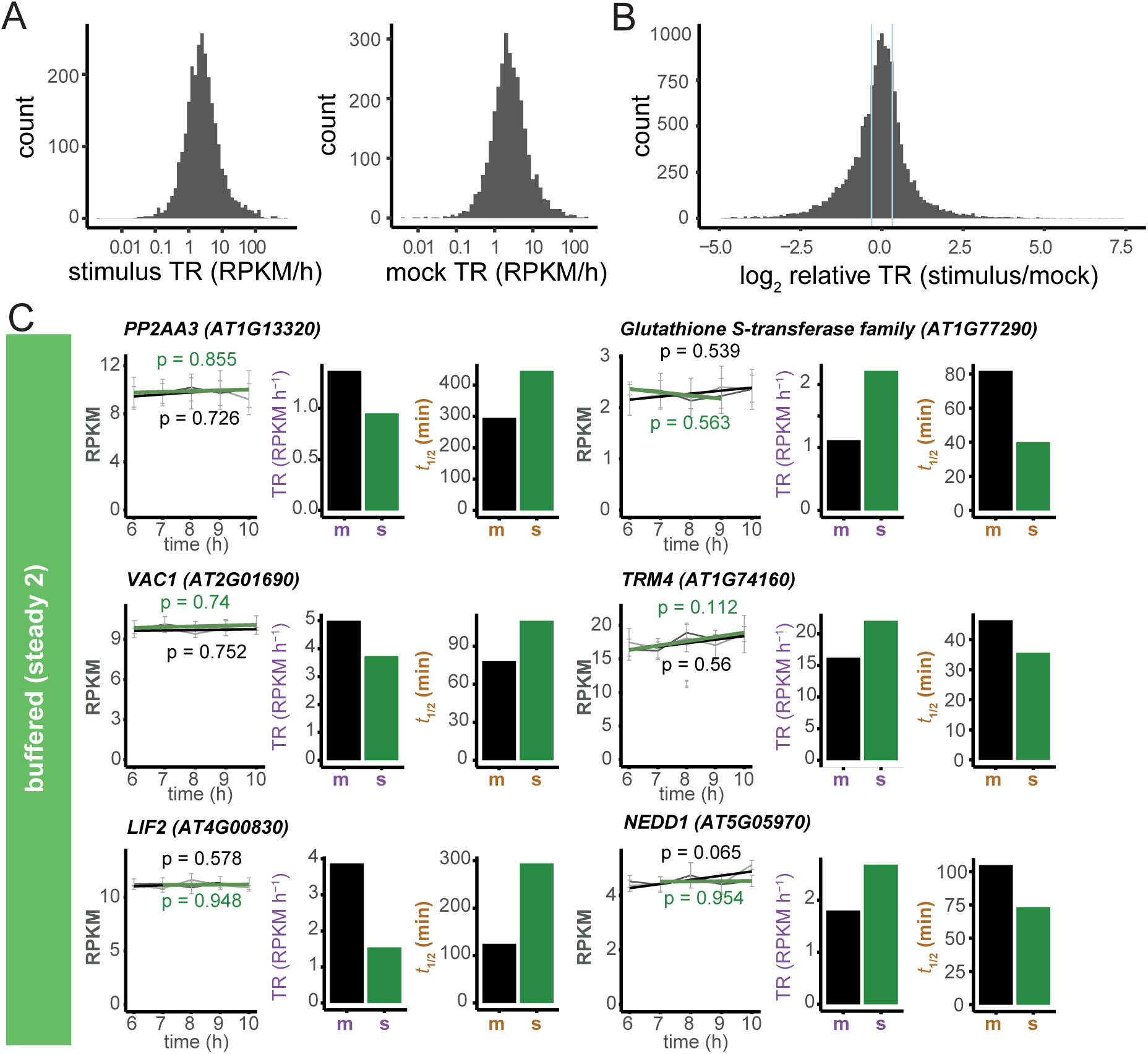
Early stimulus transcriptional response is varied and dynamic and includes buffered RNAs. (A) Distributions of TRs for mock and stimulus RNAs with changing abundances presented as histograms. (B) Distribution of log2 relative TRs (stimulus/mock) as a histogram. Solid light blue lines indicate the cutoffs for unchanged TR (1 +/- 0.18). (C) Examples of RNA buffering from Steady group 4. Buffered RNAs have steady abundances but their *t*1/2s and TRs differ. For each example RNA, abundance measurements (RPKM mean±SE) at 6, 7, 8, 9, 10 h of mock (thin light gray line) and stimulus (thin dark gray line) along with linear models of rate of abundance change are shown for mock (thick black line) and stimulus (thick green line). Model p-values label the graphs. *t*1/2s and TRs are also shown in bar plots to the right for mock (m), and stimulus (s).

**Supplemental Figure S5.**
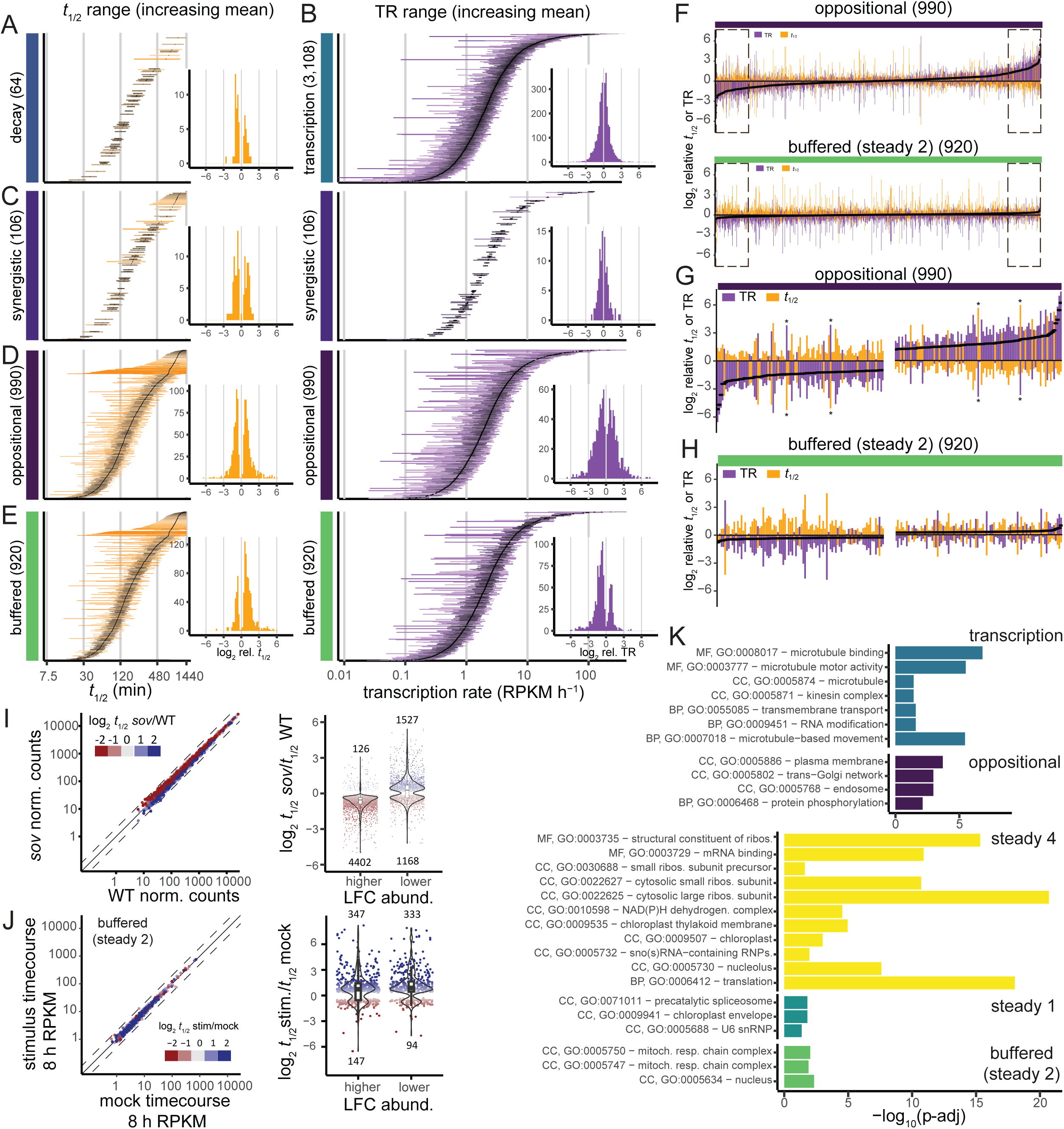
Dynamic range extended analysis. (A,C-E) Dynamic range of decay rate shifts (orange line segments, left column) in regulatory modalities. Line segment endpoints indicate mock and stimulus *t*1/2s. RNAs are ordered on the y-axis by increasing *t*1/2 mean of mock and stimulus treatments (indicated by black circles). Inset histograms show distribution of log2 relative *t*1/2. (B,C-E) Dynamic range of TR changes (purple line segments, right column) in regulatory modalities. Line segment endpoints indicate mock and stimulus TRs. RNAs are ordered on the y-axis by increasing mean TR of mock and stimulus and treatments (indicated by black circles). Inset histograms show distribution of log2 relative TR. (F) Oppositional and Steady 2: Buffered stacked bar plots with dashed-line boxes indicating expanded regions in (G) and (H), respectively. Asterisks highlight exceptionally large and nearly symmetric changes in TR and *t*1/2. (I,J) Comparison of abundances and *t*1/2 shifts of RNA buffered in *sov* mutants and buffered (Steady 2). (K) Gene ontology enrichment analysis of regulatory modality RNAs. The decay, synergistic, and steady 3 modalities did not have enriched GOs (p-adj, adjusted p-value).

